# Biventricular interaction during acute left ventricular ischemia in mice: a combined in-vivo and in-silico approach

**DOI:** 10.1101/2023.01.26.525736

**Authors:** M. J. Colebank, R. Taylor, T. A. Hacker, N.C. Chesler

**Affiliations:** Edwards Lifesciences Foundation Cardiovascular Innovation and Research Center, and Department of Biomedical Engineering, University of California, Irvine, Irvine, CA, USA; Cardiovascular Research Center, University of Wisconsin-Madison, Madison, WI, USA

**Keywords:** Computational model, parameter estimation, myocardial infarction, biventricular interaction, sensitivity analysis, multiscale modeling

## Abstract

Computational models provide an efficient paradigm for integrating and linking multiple spatial and temporal scales. However, these models are difficult to parameterize and match to experimental data. Recent advances in both data collection and model analyses have helped overcome this limitation. Here, we combine a multiscale, biventricular interaction model with mouse data before and after left ventricular (LV) ischemia. Sensitivity analyses are used to identify the most influential parameters on pressure and volume predictions. The subset of influential model parameters are calibrated to biventricular pressure-volume loop data (n=3) at baseline. Each mouse underwent left anterior descending coronary artery ligation, during which changes in fractional shortening and RV pressure-volume dynamics were recorded. Using the calibrated model, we simulate acute LV ischemia and contrast outputs at baseline and in simulated ischemia. Our baseline simulations align with the LV and RV data, and our predictions during ischemia complement recorded RV data and prior studies on LV function during myocardial infarction. We show that a model with both biventricular mechanical interaction and systems level cardiovascular dynamics can quantitatively reproduce *in-vivo* data and qualitatively match prior findings from animal studies on LV ischemia.

## 1. Introduction

Coronary artery disease, which leads to myocardial infarction, accounts for roughly 41% of all cardiovascular-related deaths^30^. Acutely disrupted blood flow and oxygen supply to the myocardium causes cell death and systolic dysfunction, raising diastolic ventricular and atrial filling volumes^2^. Increases in left ventricular (LV) volume raise left atrial and pulmonary venous pressure^8^, the latter of which is hypothesized to initiate vascular remodeling and pulmonary hypertension with the eventual consequence of right heart failure^1,23^. This cascade of events is difficult to integrate from experimental or clinical data alone, yet a better understanding of the acute effects of LV ischemia will provide insight into long-term cardiac and vascular remodeling. Hence, here we combine an *in-silico* computational model of biventricular interaction with *in-vivo* data from a cohort of male mice subjected to LV ischemia.

The right ventricle (RV) is mechanically linked to the LV through the interventricular septum (S). Previous canine studies^6^ in the absence of RV electrical pacing reported that 68% of RV systolic pressure and 80% of pulmonary flow output were attributed to LV and S contributions. Follow up investigations^11^ also reported that RV ischemia reduced pulmonary systolic pressures by 4 mmHg, while septal ischemia had a greater effect on the RV and reduced pulmonary systolic pressures by 8 mmHg. Thus, systolic dysfunction in either chamber impairs function, which emphasizes the importance of biventricular interaction in cardiac function.

*In-vivo* experiments usually provide insightful but isolated measurements of cardiovascular function. Integration of this data can deliver new information regarding the underlying physiological mechanisms. *In-silico* computational models are a promising tool for integrating multimodal data from *in-vivo* experiments and testing mechanistic hypotheses surrounding disease progression. For example, early work using isolated ventricular elastance models in a closed loop compartment model investigated the link between LV systolic dysfunction and pulmonary venous pressure^3^. While reduced LV end-systolic elastance alone could not replicate the rise in pulmonary venous pressure seen clinically, additional increased systemic venous volume and pericardial constraints in the model framework could recreate these established findings. Efforts have also resulted in the incorporation of LV remodeling and hemodynamic reflexes^34^, which synergistically contribute to LV remodeling.

These prior computational studies did not explicitly account for biventricular interaction or include multiscale mechanisms. The cutting-edge reduced order model of ventricular interaction is the three-segment (“TriSeg”) model by Lumens et al., which represents the LV, RV, and S as thick walled, spherical chambers driven by myocyte dynamics^16^. Several authors have had success in using this framework to simulate disease, such as pulmonary hypertension^29^ and LV ischemia^15^. These models contain numerous parameters, requiring a formal model analysis to determine which parameters are influential and identifiable given limited data^4^. The combination of multiscale *in-silico* models of biventricular interaction with robust parameter estimates from *in-vivo* data provide a necessary tool in linking experimental measurements to cardiovascular function.

Here, we combine our previously reported multiscale model^4^ with data from a cohort of male mice in baseline and acutely ischemic conditions. Echocardiographic, pressure, and volume data from the LV, RV, and systemic arteries are collected pre-ischemia. We determine a subset of influential parameters using sensitivity analyses and calibrate the multiscale model to baseline data. Our model predictions at baseline align with the measured data across all animals. We simulate acute ischemia by reducing LV active force, and report increased LV end-diastolic volumes, reduced LV longitudinal strain, and elevated left atrial pressure. Continuous recordings of RV pressure-volume loops during acute ischemia are contrasted to the model predictions. Lastly, we compare simulated LV pressure versus sarcomere length, which qualitatively agree with previous findings^17^.

## 2. Methods

### 2.1 *In-vivo* animal data

All animal procedures were approved by the University of Wisconsin-Madison Institutional Animal Care and Use Committee. Three adult C57/B16 male mice (20-22 weeks old) were anaesthetized with 5% isoflurane and maintained with 1-2% isoflurane and room air throughout all procedures. Mice were put on a heated platform to maintain a body temperature of 37°C and measure ECG activity. Transthoracic echocardiography (Vevo 3100, Visual Sonics) was used to identify systolic and diastolic inner diameter and fractional shortening for both the LV and RV. A cutdown was performed on the right carotid artery and a 1.2 Fr pressure catheter (Transonic) was placed and advanced to the ascending aorta to measure systemic pressures. Finally, the thoracic cavity was entered, and the heart was exposed. A 1.2 Fr pressure-volume catheter with 4.5 mm spacing (Transonic) was inserted into the LV via direct stick through the myocardial wall. Baseline systemic and LV data was recorded. The catheter was removed and a second 1.2 Fr pressure-volume catheter with 3.5mm spacing (Transonic) was place in the RV free wall aligned with the pulmonary valve. Baseline systemic and RV data were collected. A 7-0 suture was placed around the left anterior coronary artery mid ventricle and tied while still recording RV data. Typical ECG changes and blanching were noted. Pressure and volume measurements were recorded at 500 Hz and analyzed on commercially available software (Notocord Systems, Croissy Sur Seine, France). After, the mice were sacrificed and the four heart chambers were dissected and weighed^23^. Heart chamber weight is converted to wall volume using a constant density of 1.053 g/ml and used in the computational model described later. We assume the septum occupies 1/3 of the LV volume^19^. A schematic of the experimental design is provided in Figure 1(a).

**FIGURE 1:**
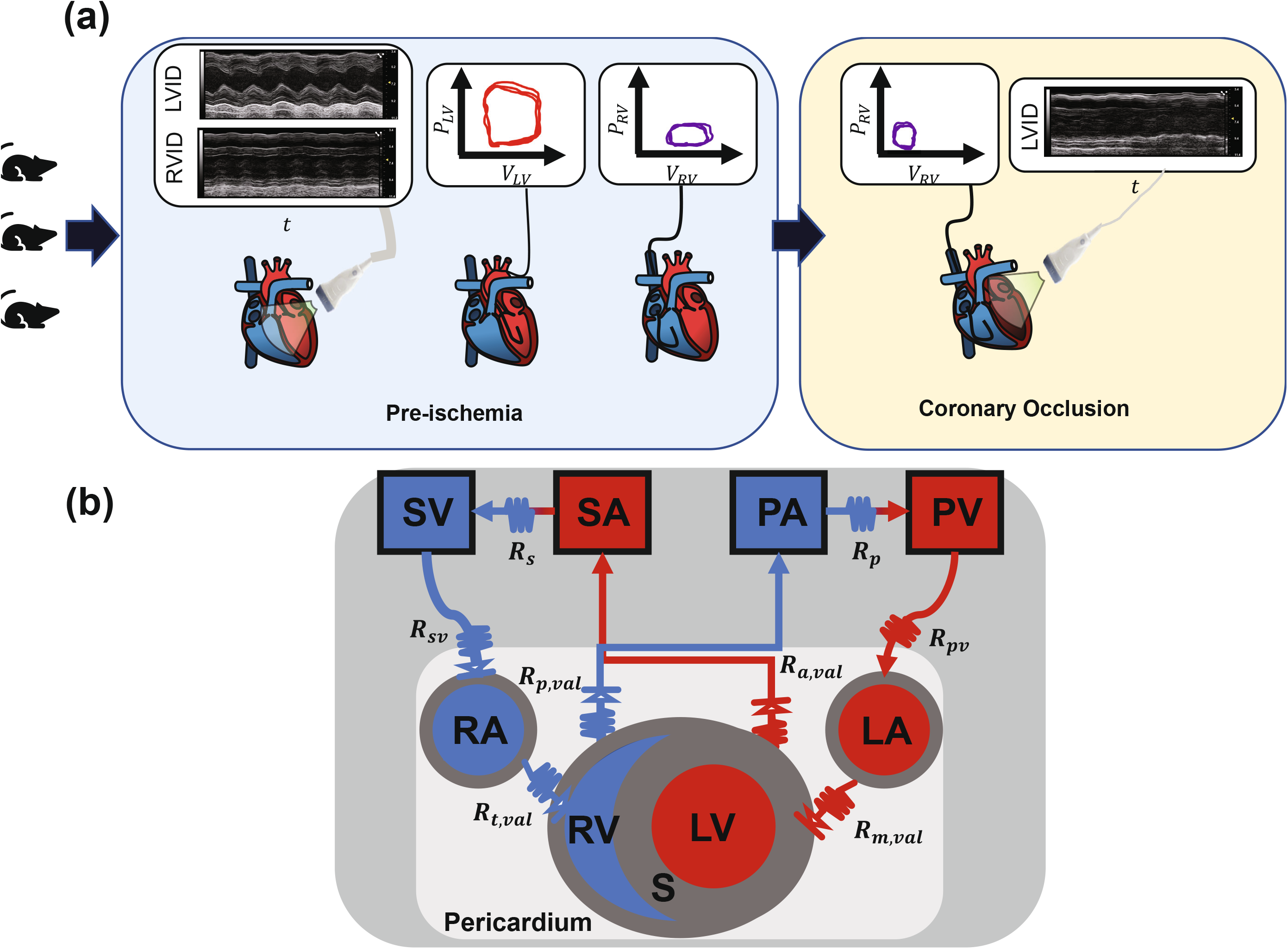
Experimental and model schematics. (a) Three male mice underwent non-invasive echocardiography, providing measurements of ventricular inner diameter. A pressure-volume catheter was then placed in the LV chamber, data were recorded, the catheter was removed, and placed in the RV. While the RV catheter was still in, the left anterior descending coronary artery was ligated, and RV pressure-volume data was recorded. Echocardiography was repeated. (b) Schematic of the closed loop computational model. The two ventricles are coupled through a dynamic septal wall using the TriSeg framework. All four heart chambers are encased in a passive, pericardial sack and connected to compliant arterials and venous compartments. Resistors connect all model components. LA – left atrium; LV – left ventricle; PA – pulmonary arteries; PV – pulmonary veins; RA – right atrium; RV – right ventricle; S – septum; SA – systemic arteries; SV – systemic veins.

The built in Gaussian smoothing filter in MATLAB (Mathworks, Natick, MA) was used to remove noise in the pressure-volume signals. In house algorithms were used to separate signals into beat-by-beat datasets for analyses. To account for discrepancies in pressure-volume phase due to catheter placement, volume traces were slightly shifted to ensure maximal chamber volume occurred at the upstroke of ventricular pressure.

### 2.2 Mathematical model

We use a previously developed multiscale cardiovascular model^15,16^. The model components include 1) a modified Hill model of sarcomere shortening, 2) an empirical model of cardiomyocyte calcium handling, 3) four spherical cardiac chambers including biventricular interaction, and 4) a zero-dimensional (0D) hemodynamics model.

The sarcomere length *L_s_* (*μ*m) is determined from the myofiber strain, *ε_f_* within each chamber

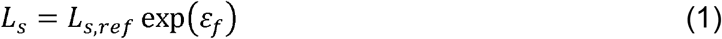

where *L_s,re1_* = 0 2.0 (*μ*m) is the reference sarcomere length at zero strain (i.e.,*ε_f_* = 0). The contractile sarcomere element has length *L_sc_* (*μ*m) and is in series with an elastic series element with length *L_se_* = *L_s_*−*L_sc_* (*μ*m). Sarcomere shortening is described by

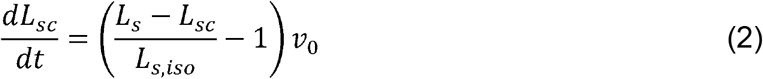

where *L_s,iso_* (*μ*m) is the elastic series element length in an isometrically stressed state, and *v_o_* (*μ*m/s) is the velocity of sarcomere shortening with zero load^16^. Sarcomere activation is modeled as the sum of a rise and decay terms

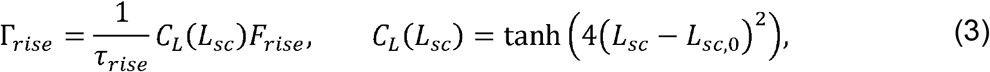

where *C_L_* (dimensionless) represents the increase in contractility with sarcomere length and *L_sc,o_* (*μ*m) represents the contractile element length with zero active stress. The second term *F_rise_* (dimensionless) describes changes in cardiomyocyte intracellular calcium

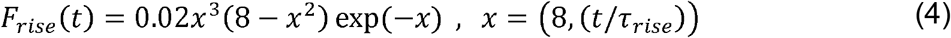

where *τ_rise_* (s) scales the rise in contractility. Calcium decay is given by

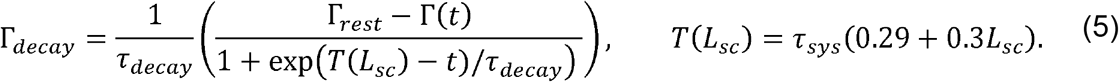

The decay in activation saturates at the diastolic value Γ_rest_ (dimensionless), and depends on the systolic contraction and diastolic decay parameters, *τ_sys_* and *τ_decay_* (s), respectively. Equations (3) and (5) dictate the contractile differential equation:

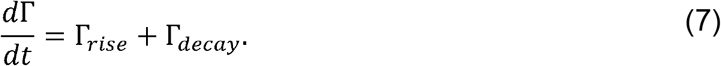

The active stress, *G_act_* (KPa) is finally calculated as

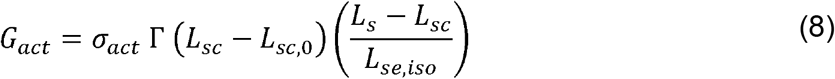

where σ_act_ (KPa) is a scaling parameter^16^. Passive sarcomere stretch is relative to the passive reference length *L_s,pas,re1_* (*μ*m)

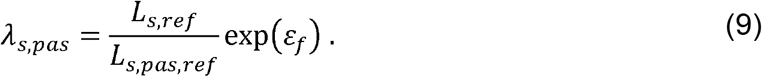

The passive stresses are separated into those attributed to the extracellular matrix (ECM) and Titin

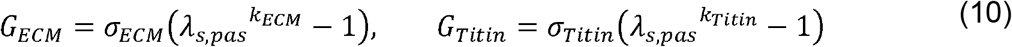

where *σ_ECM_* and *σ_Titin_* (KPa) are scaling parameters and *k_ECM_* and *k_Titin_* (dimensionless) account for nonlinear chamber stiffening^31^.

The sarcomere model is embedded within each cardiac chamber and the interventricular septum. Ventricular interaction across the septal wall is prescribed using the TriSeg model^16^. Cardiac chamber geometries are modeled as spherical structures described by a mid-wall volume *V*_m_ (mm^3^), mid-wall curvature *C_m_* (mm^−1^), and mid-wall cross-sectional area *A_m_* (mm^2^) and parameterized by a reference mid-wall area, *A_m,ref_* (mm^2^), and a wall volume, *V*_wall_ (mm^3^). Tension balance across the LV, RV, and S walls are enforced by two algebraic constraints. Details regarding the chamber equations can be found in the Supplementary Material. All four heart chambers are enclosed in a pericardium. We assume that the pericardial sack has a reference volume, *V_o,peri_*, and exhibits a nonlinear pressure-volume relationship driven by total blood volume in the heart^14^. Pericardial pressure, *p_peri_* (KPa), is then

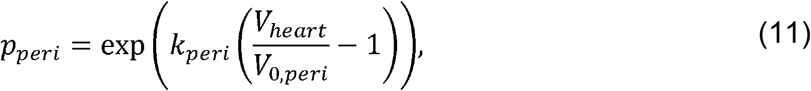

where *V_heart_* (*μ*l) represents the total volume in all four heart chambers and *K_peri_* (KPa) describes the exponential rise in pericardial pressure. This pressure value is added to each cardiac chamber as an external pressure source.

Arteries and veins are modeled as compliant compartments. Changes in blood volume *V* (*μ*l), flow *q* (*μ*l/s), and pressure *p* (KPa) are described as^5^

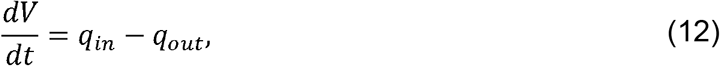

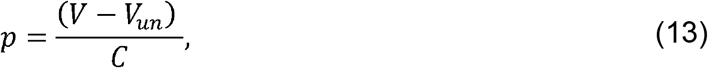

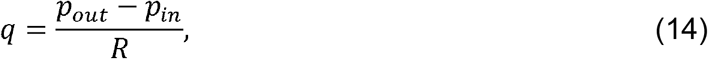

where *V_un_* (*μ*l) is the unstressed volume *C* (*μ*l KPa^−1^) is the vascular compliance, and *R* (KPa s *μ*l^−1^) is the vascular resistance between compartments. Cardiac valves are modeled as diodes and are only open when the pressure gradients are positive. An additional systemic venous valve is also included. A schematic of all model components can be found in Figure 1(b).

### 2.3 Model analysis

The mathematical model includes 18 differential equations (eight compartment volumes, *V(t)*, five sarcomere states, *L_sc_(t)*, and five contractility states, *Γ(t)*) as well as two equilibria constraints (tension balance for the TriSeg model, see Supplemental Material). These equations require a total of 53 parameters, described in Table 1, which cannot be inferred simultaneously. We fix several parameters based on prior work^4^ or available data. This leaves 38 free parameters to analyze by Morris screening and local sensitivity analysis^4,21^. Table 1 and the Supplementary Material describe how nominal parameters are calculated.

**Table 1.** Model parameters. Parameters denoted with an * are fixed before performing ventricular component, with atrial values provided in parenthesis. Mouse specific values are given in square brackets. Pressure and volume variables are described in detail in the Supplementary Material. Cap – Capillary; CO – cardiac output; LA – left atrium; LV – left ventricle; PA – pulmonary arteries; PV – pulmonary veins; RA – right atrium; RV – right ventricle; S – septum; SA – systemic arteries; SV – systemic veins.

| Parameter | Description | Units | Nominal Value/Equation |
| --- | --- | --- | --- |
| **<u>Sarcomere Parameters</u>** |  |  |  |
| $L_{s,ref}^*$ | Reference sarcomere length at zero strain | $\mu\text{m}$ | 2.0 |
| $L_{s,iso}^*$ | Elastic series element length in isometric state | $\mu\text{m}$ | 0.04 |
| $L_{s,pas,ref}^*$ | Reference length for passive wall constituents | $\mu\text{m}$ | 1.8 |
| $v_{0,j}$ | Velocity of sarcomere shortening | $\mu\text{m/s}$ | (24), 12 |
| $L_{sc,0}^*$ | Contractile element length | $\mu\text{m}$ | 1.51 |
| $\Gamma_{rest}^*$ | Resting contractility | Dimensionless | 0.02 |
| $\tau_{rise,j}$ | Rise in contractility scaling | s | $(0.0375 \cdot T)$, 0.009 |
| $\tau_{decay,j}$ | Decay in contractility scaling | s | $(0.005 \cdot T)$, 0.009 |
| $\tau_{sys,j}$ | Length of systole | s | $(0.15 \cdot T)$, 0.038 |
| $\tau_{offset,A}$ | Offset of atrial systole | s | $0.18 \cdot T$ |
| $k_{ECM,j}$ | Nonlinear ECM stiffness exponent | Dimensionless | (10), 10 |
| $k_{Titin,j}$ | Nonlinear Titin stiffness exponent | Dimensionless | (6), 6 |
| $\sigma_{ECM,j}$ | Passive ECM stress scaling factor | KPa | (0.08), 0.08 |
| $\sigma_{Titin,j}$ | Passive Titin stress scaling factor | KPa | (0.25), 0.25 |
| $\sigma_{act,j}$ | Active stress scaling factor | KPa | (35), 75 |
| **<u>TriSeg/Cardiac Parameters</u>** |  |  |  |
| $V_{LA,wall}^*$ | LA wall volume | mm<sup>3</sup> | Chamber Weight × 1.053g/mL |
| $V_{LV,wall}^*$ | LV wall volume | mm<sup>3</sup> | Chamber Weight × 1.053g/mL |
| $V_{RA,wall}^*$ | RA wall volume | mm<sup>3</sup> | Chamber Weight × 1.053g/mL |
| $V_{RV,wall}^*$ | RV wall volume | mm<sup>3</sup> | Chamber Weight × 1.053g/mL |
| $V_{S,wall}^*$ | S wall volume | mm<sup>3</sup> | Chamber Weight × 1.053g/mL |
| $A_{m,ref,LA}$ | LA reference area | mm<sup>2</sup> | [0.15,0.20,0.20] |
| $A_{m,ref,LV}$ | LV reference area | mm<sup>2</sup> | [0.50,0.70,0.50] |
| $A_{m,ref,RA}$ | RA reference area | mm<sup>2</sup> | [0.15,0.20,0.20] |
| $A_{m,ref,RV}$ | RV reference area | mm<sup>2</sup> | [0.55,0.80,0.50] |
| $A_{m,ref,S}$ | S reference area | mm<sup>2</sup> | [0.25,0.35,0.35] |
| $V_{0,peri}$ | Reference volume for pericardial space | μL | [0.159,0.178,0.165] |
| $k_{peri}$ | Material parameter of pericardial tissue | KPa | $10^{14}$ |
| **<u>Cardiovascular System Parameters</u>** |  |  |  |
| $R_{a,val}$ | Aortic valve resistance | KPa s/ μL | $0.0133 \div CO$ |
| $R_{m,val}$ | Mitral valve resistance | KPa s/ μL | $0.067 \div CO$ |
| $R_{p,val}$ | Pulmonic valve resistance | KPa s/ μL | $0.013 \div CO$ |
| $R_{t, val}$ | Tricuspid valve resistance | KPa s/ μL | $0.067 \div CO$ |
| $R_{vc}$ | Vena Cava resistance | KPa s/ μL | $(\bar{P}_{sv} - P_{ra, min}) \div CO$ |
| $R_{pv}$ | Pulmonary venous resistance | KPa s/ μL | $(\bar{P}_{pv} - P_{la, min}) \div CO$ |
| $R_{sys}$ | Systemic circulation resistance | KPa s/ μL | $(P_{sa, max} - P_{cap, sys'}) \div CO$ |
| $R_{pulm}$ | Pulmonary circulation resistance | KPa s/ μL | $(P_{pa, max} - P_{cap, pulm}) \div CO$ |
| $C_{sa}$ | Compliance of systemic arteries | μL / KPa | $V_{sa} \div P_{sa, max}$ |
| $C_{sv}$ | Compliance of systemic veins | μL / KPa | $V_{sv} \div \bar{P}_{sv}$ |
| $C_{pa}$ | Compliance of pulmonary arteries | μL / KPa | $V_{pa} \div P_{pa, max}$ |
| $C_{pv}$ | Compliance of pulmonary veins | μL / KPa | $V_{pv} \div \bar{P}_{pv}$ |

Morris screening is an efficient screening tool that uses coarse approximations of model sensitivity to determine which parameters are non-influential^21^. We use model predictions LV and RV pressure-volume relationships as well as systemic arterial pressure as our quantities of interests. We rank parameter importance based on the modified sample mean, μ^*^ and sample variance, *S*^2^, through the index 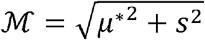 ^4,33^ Similar to van Osta et al^21^, parameters consistently less influential than the mean value of 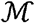 on all five outputs are deemed non-influential and fixed. Parameter bounds are set at ±20% from the nominal value for each mouse.

Local sensitivity analysis is performed around the nominal values of the remaining parameters. The local sensitivity of each pressure or volume, denoted as *F(t;**θ***), is approximated by centered finite differences

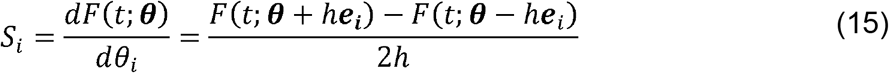

where *h* = 0.01 is the step size and *e_i_* is the unit vector in the i-th direction. To account for differences in magnitudes we use dimensionless sensitivities by multiplying by *θ_i_/F(t;**θ***)^20^. We use the local sensitivity vectors to construct an approximate Fisher information matrix, ***F** = **s^τ^s***, and assess practical identifiability in an asymptotic sense^5,9^. If **F** is ill-conditioned, then the parameter subset is deemed non-identifiable the matrix is no longer ill-conditioned. We iterate this scheme until cond(***F***) ≤ 10^5^, is our numerical ill-conditioning cutoff.

### 2.4 Parameter inference and uncertainty quantification

The reduced subset is calibrated to data using nonlinear least squares^5^. We combine LV, RV, and aortic data into a single residual vector

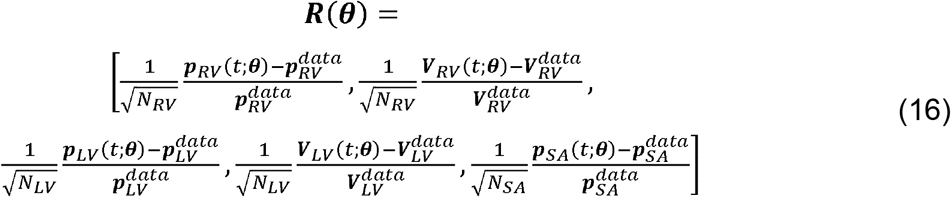

with the superscript `data` denoting the measured data, and *N_Rv_, N_Lv_*, and *N_sA_* representing the number of data points within the RV, LV, and aortic time series signals, respectively. Mouse specific parameters 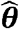 are using the built in *lsqnonlin* function in MATLAB.

We quantify parametric and output uncertainty after determining the optimal parameter values. Let 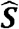 denote the sensitivity vector at 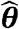 for each mouse. Using the minimized residual, 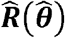, the parameter confidence intervals are^27^

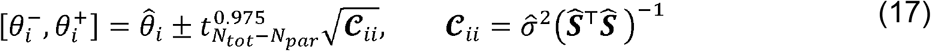

where 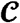 is the asymptotic sample covariance matrix, 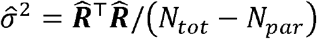 is the sample noise variance using the number of total data points *N_tot_* and the number of parameters N_par_, and 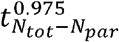 is a two-sided t-score statistic corresponding to a 95% confidence interval^27^. Corresponding confidence and prediction intervals for the optimal model output, 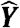, are

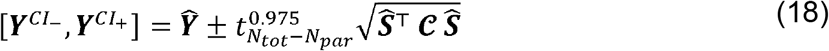

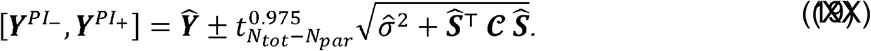

Though the data spans multiple heart beats, the model does not change from beat to beat, hence 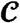 is constructed using a single model cycle.

### 2.5 Simulated Myocardial Infarction

We simulate myocardial infarction by reducing LV active force at the sarcomere

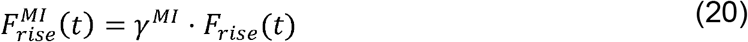

where γ^*MI*^ reflects the decrease in activation due to ischemia. We set γ^*MI*^ such that LV ejection fraction is reduced by the same amount measured by echocardiography. We also examine changes in longitudinal wall strain

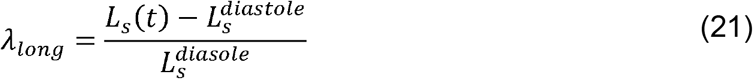

where *L_s_(t)* is the dynamic sarcomere length and 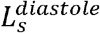 is the length at end-diastole.

## 3. Results

### 3.1 *In-vivo* data

Echocardiography and pressure-volume loops for each mouse are shown in Figure 2. The time dependent ventricular pressure and volume data are provided for each mouse, including RV pressure-volume data during ischemia in Figure 2(c). Ischemia introduces a decrease in RV volumes, especially in mouse 3. LV and RV inner diameters (Figure 2(d)) are similar across all three mice. After coronary artery ligation, there is an increase in both systolic and diastolic LV inner diameter, contributing to a reduction in fractional shortening. There is also an increase in RV diastolic diameter, but not in RV systolic diameter nor in fractional shortening.

**FIGURE 2:**
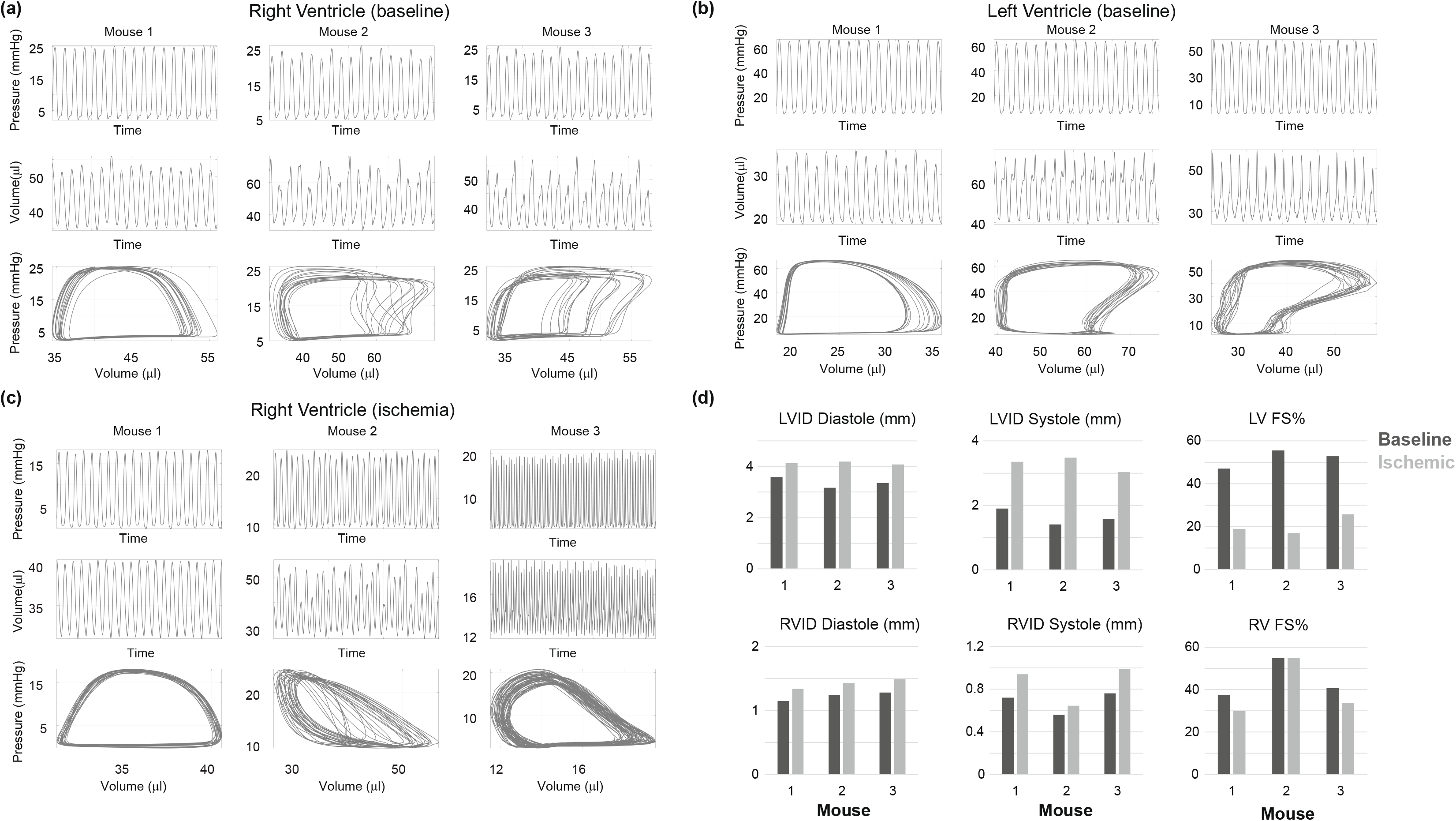
*In-vivo* data from three male mice. (a) Pressure, volume, and combined pressure-volume loops in the RV at baseline. (b) Pressure and volume data in the LV at baseline. (c) Pressure-volume data in the RV after left descending coronary artery ligation. (d) Baseline and ischemic echocardiography measurements in the LV and RV.

### 3.2 Sensitivity analyses

A total of 100 Morris screening initializations were run per mouse. Parameter ranking for the five different model outputs are provided in Figure 3. The parameters describing the timing of ventricular systole and diastole (*τ_rise,v_*,*τ_decay,v_*, and *τ_sys,v_*) are consistently the most influential on LV and RV pressure. The vascular parameters *R_sys_*,*R_pulm_*, and *C_sv_* are also influential on both pressure predictions. The LV, RV, and S reference areas are more influential on ventricular volume than ventricular pressure. Active force generation *σ_act,v_* and the reference pericardial volume *V_o,peri_* are moderately influential for all five outputs. Eighteen parameters have an average effect less than the mean, 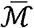, and are deemed non-influential.

**FIGURE 3:**
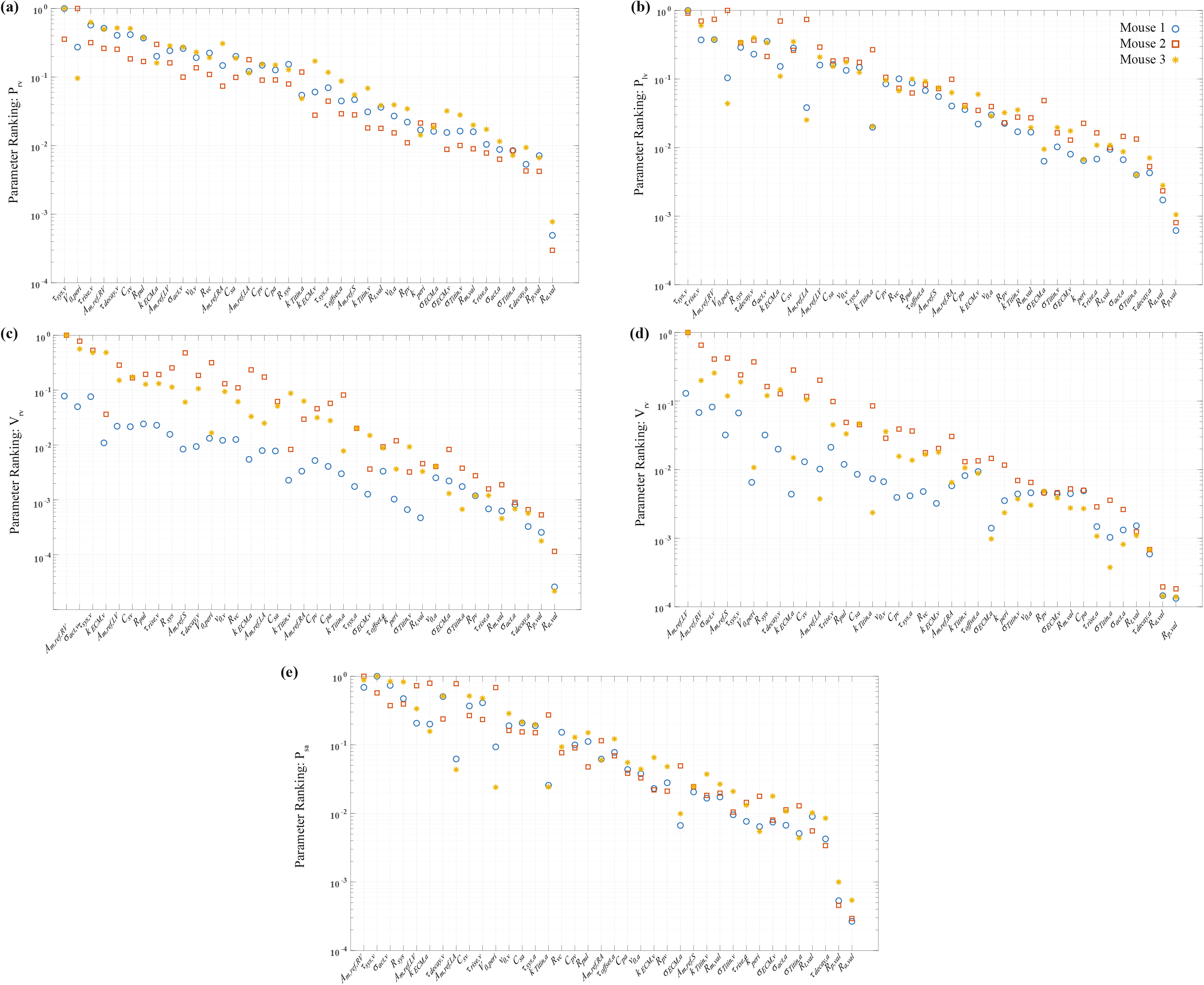
Parameter ranking using the combined index, 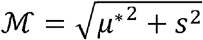, based on Morris screening for each mouse. (a) RV pressure. (b) LV pressure. (c) RV volume. (d) LV volume. (e) Systemic artery pressure. Each plot is normalized by the maximum index value for each mouse so that indices are scaled 0 to 1.

The remaining 20-parameter subset is examined using local sensitivity analysis for each mouse. The matrix ***F*** is invertible for all three mice but has a condition number iterations of subset reduction are carried out until ***F*** has condition number below 1e5. A between 1e7 to 1e8, which is numerically ill-conditioned by our criteria. Several final subset using this approach consists of the 11 parameters

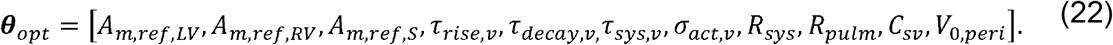

### 3.3 Model calibration and uncertainty quantification

We infer ***θ**_opt_* for each mouse using the recorded baseline data. Optimal parameter estimates and the associated confidence intervals are provided in Table 2. Simulated pressure-volume loops, shown in Figure 4(a), align well with the recorded systolic and diastolic values. However, our simulations maintain the “ideal” pressure-volume loop shape while the data does not. Confidence and predictions intervals for the time-series model outputs are shown in Figure 4(b). While beat-to-beat differences in the data are not captured by the model, confidence and predictions intervals contain nearly all the data across every heartbeat. Model simulations of pressure match well to the data, while the model predictions of volume show a slight discrepancy during isovolumic contraction.

**Table 2:**
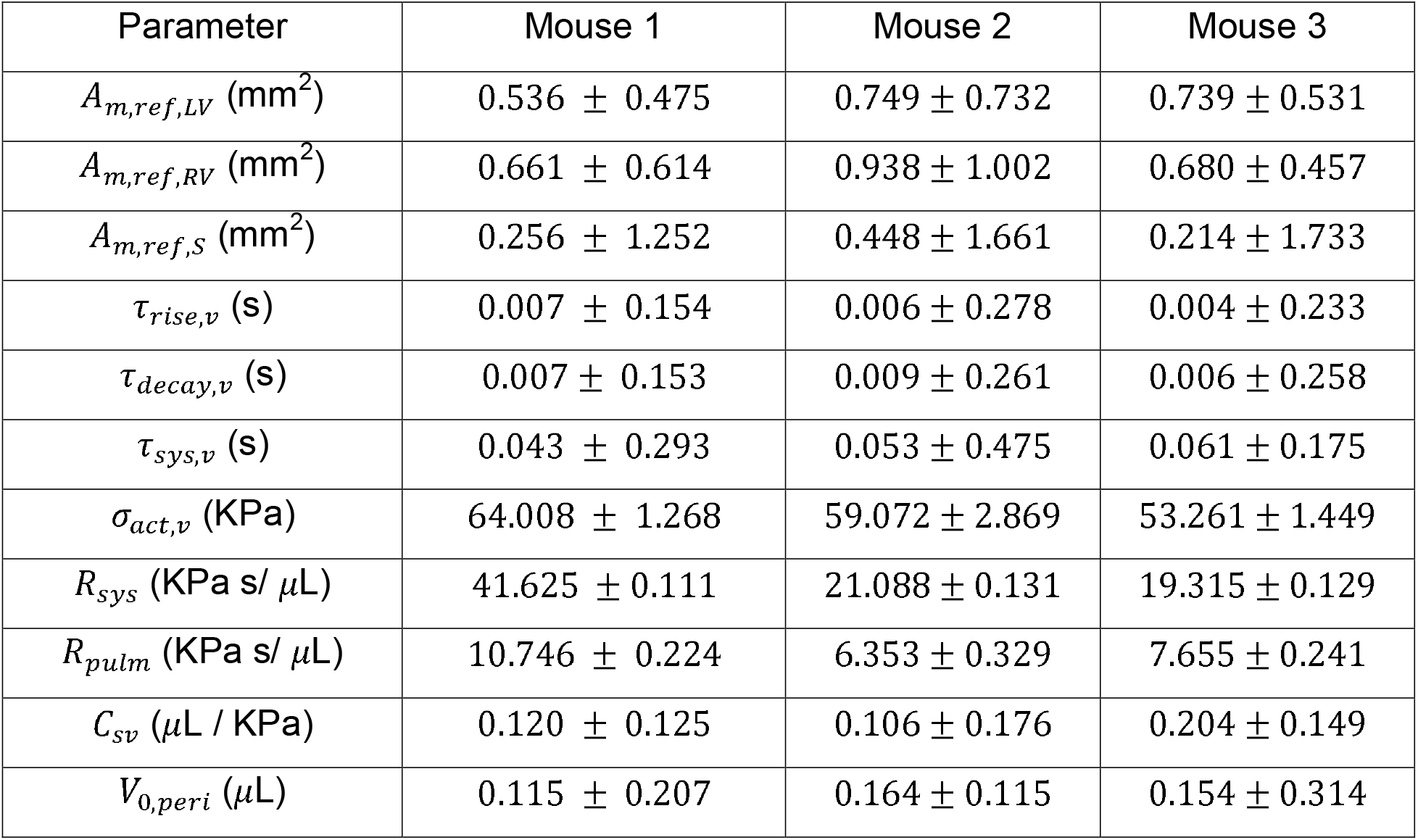
Optimal parameter estimates and “ one standard deviation.

**FIGURE 4:**
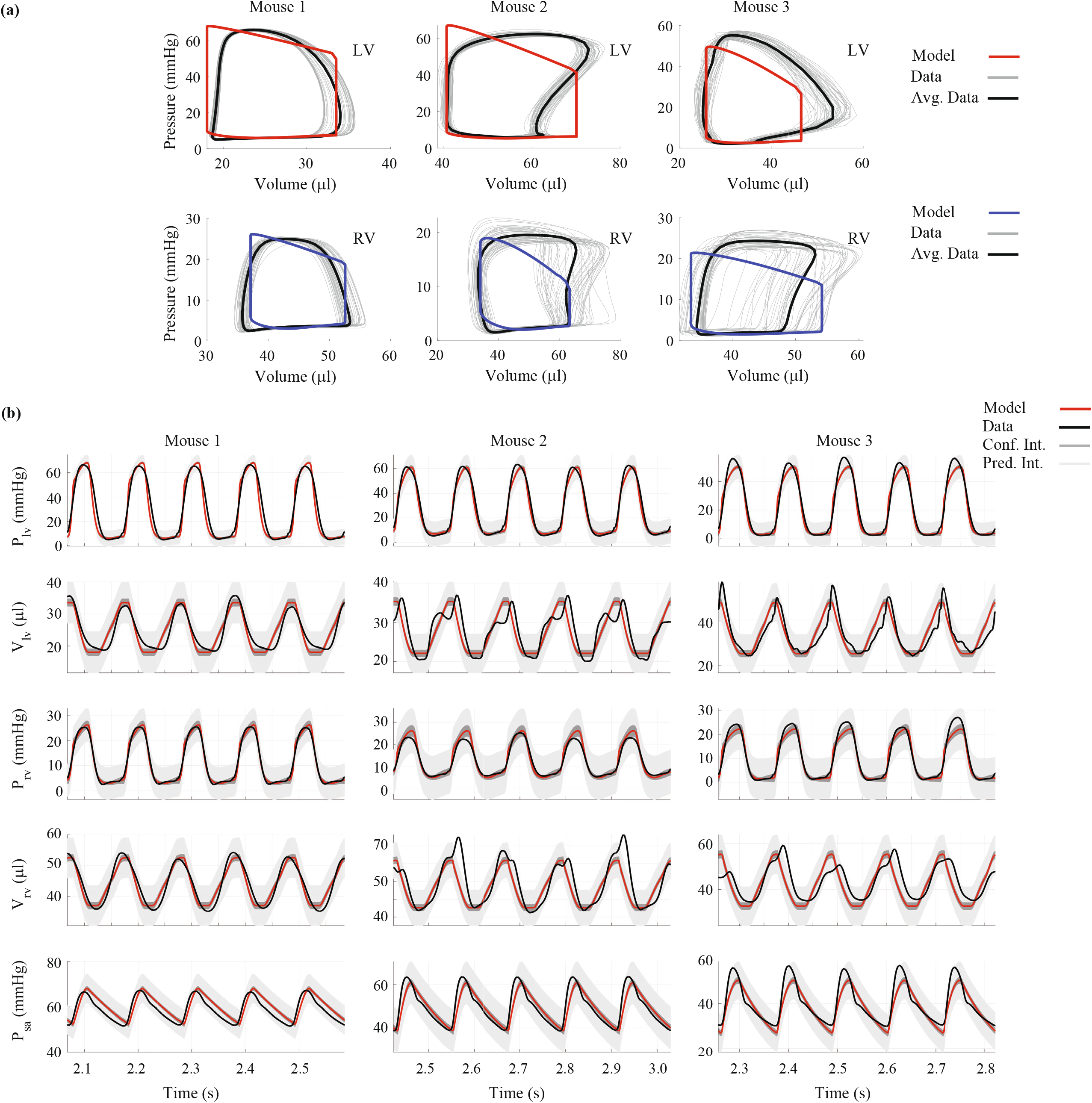
Comparison of model simulations with measured hemodynamic data. Pressure volume loops in the LV and RV (a) for each mouse. Red and blue curves represent the model predictions after parameter calibration to the measured data, shown in grey. Panel (b) shows optimal model solutions (red), confidence intervals (light grey), and predictions intervals (dark grey) for LV pressure, LV volume, RV pressure, RV volume, and SA pressure, respectively. Note that most of the beat-to-beat signals are captured within the uncertainty bounds.

### 3.4 Simulated Myocardial Infarction

We simulate LV ischemia by decreasing the active force generation using *γ^*MI*^*. *In-vivo* results in Figure 2(d) show a 50-70% reduction in EF during ischemia, (60%, 69%, and 54%, for mouse 1, 2, and 3, respectively) which is our target for the model. Simulated LV and RV outputs at baseline and in ischemia are shown in Figure 5a. Using *γ^*MI*^* = 0.20 reduced the ejection fraction by 61%, 62%, and 54%, for mouse 1, 2, and 3, respectively. Decreased contractile function shifts LV pressure-volume relationships rightward. Stroke volume in both heart chambers is reduced in ischemia, while diastolic RV pressure increases. Recorded RV pressure-volume data during ischemia also shows a slight rightward shift as seen by the model. Data from mice 1 and 2 have a similar stroke volume to that predicted by the model, while mouse 3 has a substantial reduction in volume values. Ventricular stroke work, the area within the pressure-volume loop, is shown in Figure 5b for both the data and model simulations. Stroke work is greater in the LV than the RV due to the difference in pressure magnitudes. Stroke work in both cardiac chambers decrease with LV ischemia.

**FIGURE 5:**
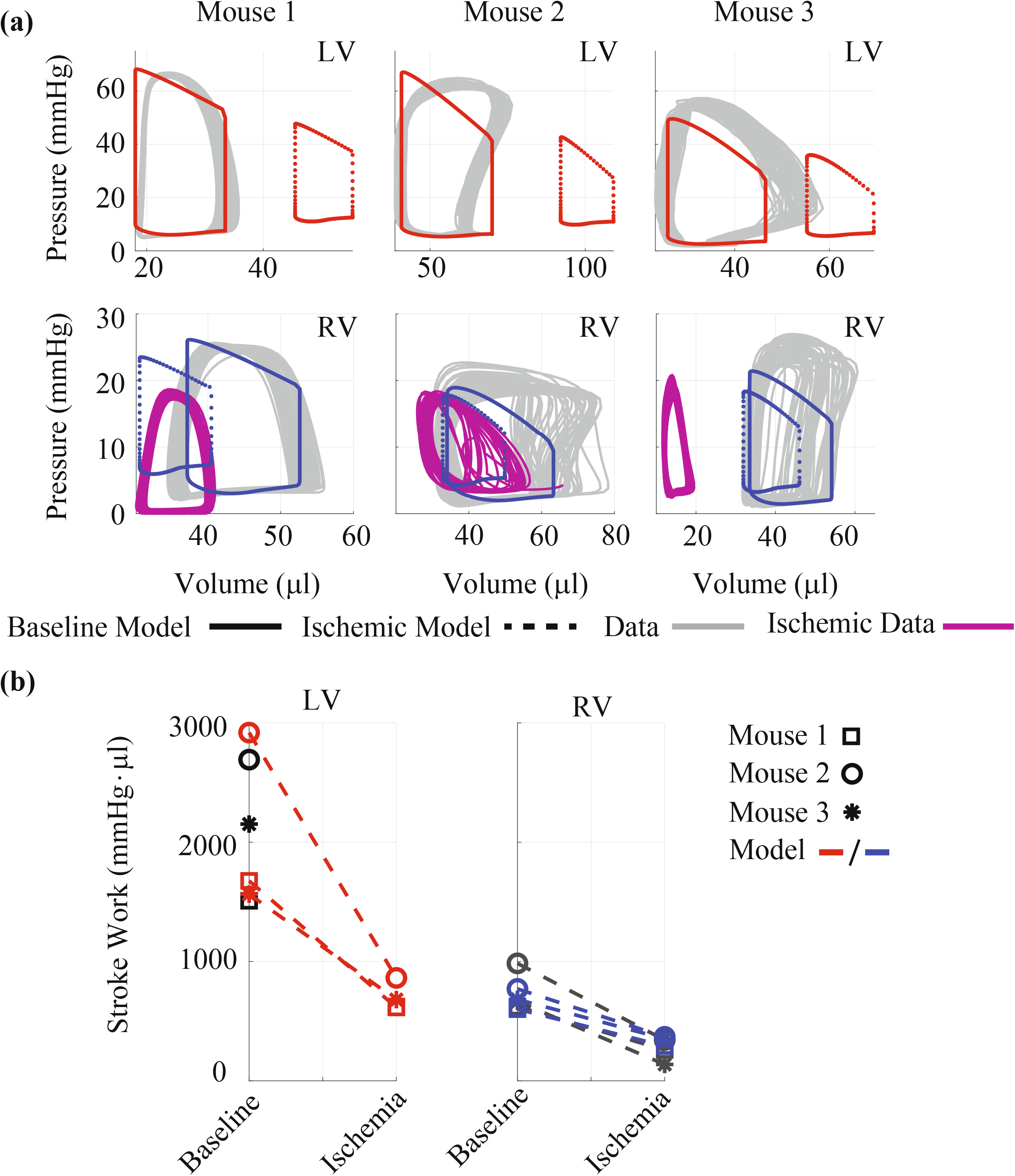
Changes in pressure-volume relationships with LV ischemia. (a) Baseline pressure-volume data (grey) in the LV (top) and RV (bottom) compared to the baseline simulations after parameter inference (solid, colored lines). Ischemic RV data (solid, blue) and RV predictions (dotted, blue). Note that LV ischemia causes a rightward shift in LV pressure-volume loops, while RV pressure-volume loops show a slight to moderate leftward shift with a reduction in stroke volume. Ischemic RV data varies with each mouse. (b) Ventricular stroke work (integral of the pressure volume loop) at baseline and in ischemia. Stroke work is larger in the LV due to pressure magnitude, and both LV and RV stroke work are reduced in ischemia.

We also investigate the systems-level effects of LV ischemia. Figure 6 displays left atrial pressure-volume loops at baseline and in ischemia. All three mice show an upward shift in ischemia, attributed to elevated LV diastolic and pericardial pressure (not shown). The latter increases on from an average 2 to 5 mmHg with ischemia.

**FIGURE 6:**
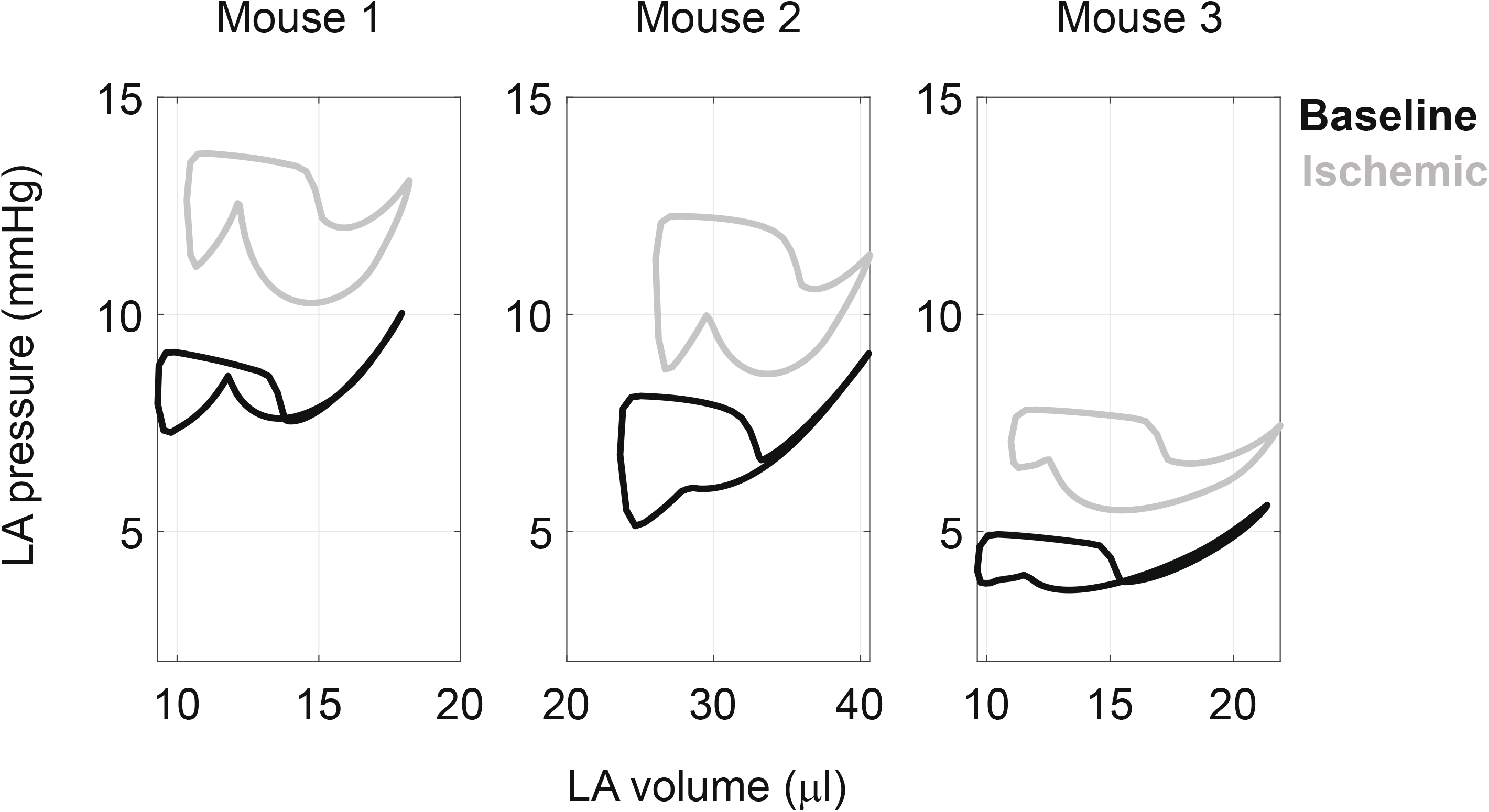
Left atrial (LA) pressure-volume loops from the model at baseline after parameter inference (black) and after LV ischemia (grey) in all three animals. The illustrated upward shift is indicative of elevated LV diastolic pressures. Note that baseline LA curves have the distinct “8” pattern seen *in-vivo*, which is absent in the ischemic case.

Longitudinal strains for the LV, RV and S at baseline and during ischemia are provided in Figure 7. Strains for all three walls are in phase at baseline, indicative of synchronous muscle shortening. In contrast, ischemic LV longitudinal strain are less pronounced due to the inability for the heart chamber to contract. RV strains are relatively unchanged with ischemia, while S wall strain magnitude is higher in systole with ischemia.

**FIGURE 7:**
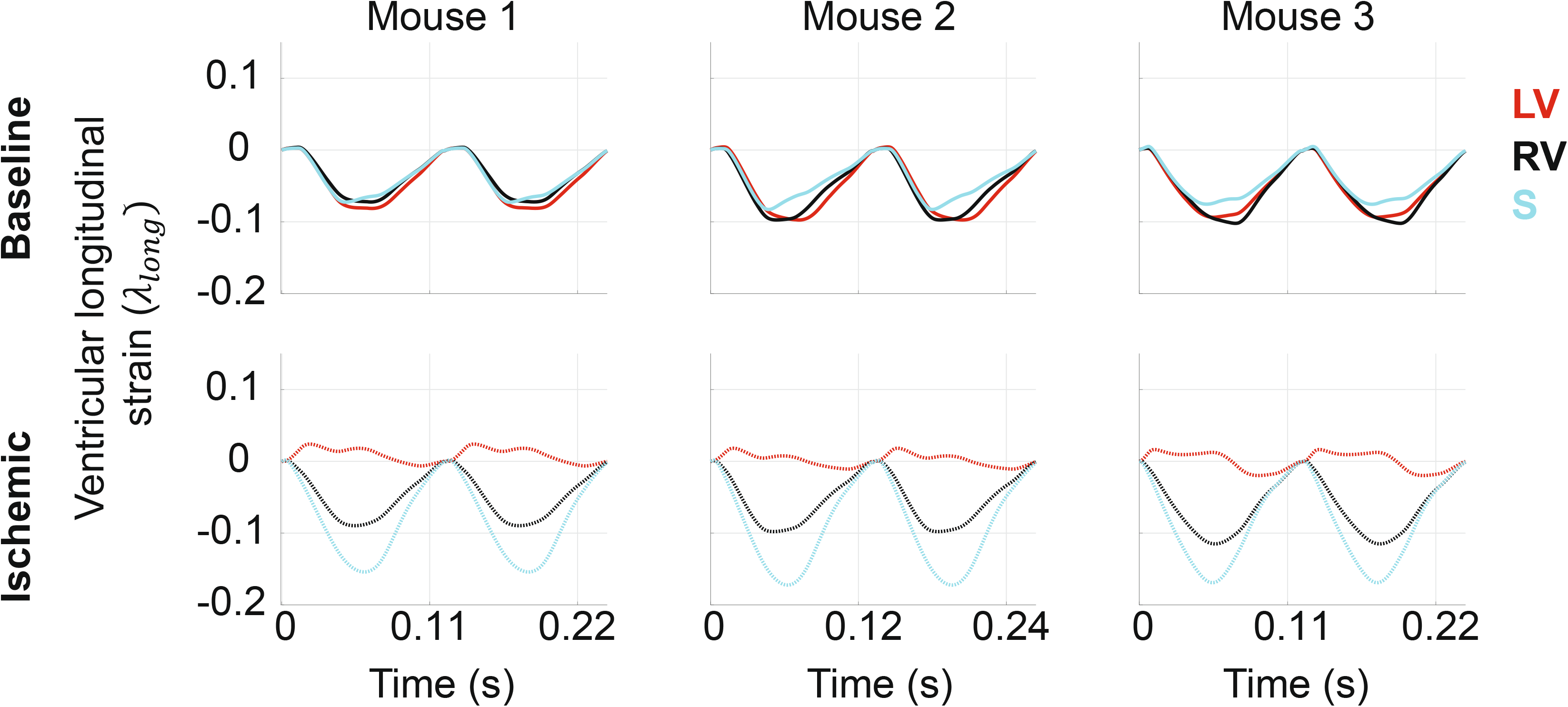
Simulated longitudinal strain in the LV, RV, and S in all three animals after parameter inference. At baseline, all three walls contract synchronously and reach 10% shortening. Ischemia reduces longitudinal strain in the LV, while RV strain is relatively unchanged and S strain is increased. Time to peak strain in the RV and S are more delayed in ischemia.

To better understand the effects of decreased LV contractility, we show LV pressure versus sarcomere length for various values of γ^*MI*^ in Figure 8. Moving from baseline (magenta, far left) to nearly aberrant active force (green, far right), results show a decrease in LV pressure and elevated sarcomere lengths. The optimal degree of reduction for the data, γ^*MI*^ = 0.2, is highlighted in the black square in Figure 8. These pressure-length curves exhibit a unique shape relative to the other pressure-length curves.

**FIGURE 8:**
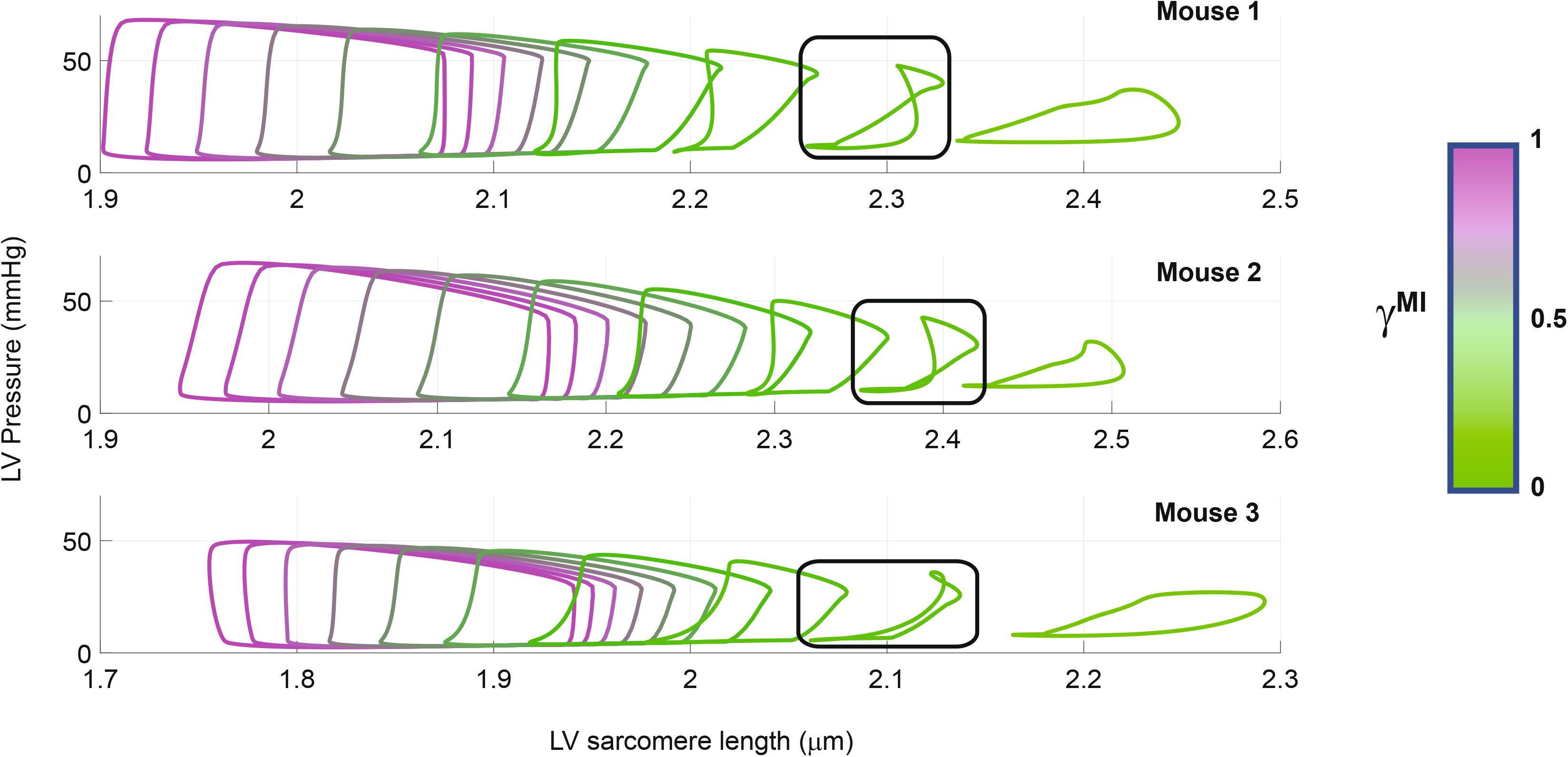
LV pressure-sarcomere length relationships in each mouse. Starting from reflecting a 0-90% decrease in LV active force generation. The value γ^*MI*^ = 0.2 is encapsulated in a black-box and provides a reduction in ejection fraction that best matches measurements in mice during ischemia. Note that the pressure-length curve has a distinct change in shape at the value of 0.2 and is qualitatively similar to prior studies using sonomicrometry^17^.

## 4. Discussion

This study combines multiscale computational modeling and parameter inference with *in-vivo* rodent data to investigate the acute biventricular consequences of LV myocardial ischemia. We identified a subset of cardiovascular parameters that can be made mouse-specific and calibrated them to hemodynamic data. We demonstrate that simulated acute LV ischemia raises septal wall strain, elevates left atrial pressure, and alters LV pressure-length relationships while RV function is relatively unchanged.

### 4.1 Model analysis

Multiscale models suffer from an imbalance in the number of model parameters versus available data. This inhibits inferring all system parameters and requires model analysis for robust parameter subsets. Morris screening is an efficient global sensitivity method to determine which parameters are non-influential, reduce simulation uncertainty, and enforce unique parameter values for each dataset^33^. Our screening results (Figure 3) show that the reference areas (*A_m,ref_*) and sarcomere contraction timing parameters (*τ_rise,v_*,*τ_decay,v_*, and *τ_sys,v_*) are consistently influential. Van Osta et al.^21^ performed a similar screening on their multiscale model of biventricular interaction. They identified reference areas, ventricular timing coefficients, and active force scaling factors as influential on ventricular wall.

We employed local sensitivity methods to further reduce our parameter space. Colunga et al.^5^ used local sensitivity to reduce their parameter subset and ensure unique, unimodal posterior distributions for Bayesian inference. Their results showed that identifiability issues arose when using a non-influential subset of parameters. The parameter subset in the current study corroborates our previous findings^4^. However, the model used here accounts for pericardial constraints. Computational studies by Pfaller et al.^22^ and Sun et al.^28^, have highlighted the importance of the pericardium on model predictions of LV and RV pressure. Sun et al. also showed that pericardial fluid volume that model predictions are sensitive to the reference pericardial volume, *v_o,peri_*.

### 4.2 Model calibration

Animal-specific computational models may provide novel physiomarkers of disease progression. These models can link data from multiple scales and multiple organs to highlight their underlying interactions. The study by Tewari et al.^29^ calibrated model parameters to match data from mice subjected to 0, 14, 21, and 28 days of pulmonary arterial hypertension conditions (using chemical and environmental stimuli)^32^. The authors found an increase in *A_m,re1,Rv_* and a decrease in *C_pa_* with increasing duration, which agrees with clinical understanding of RV and pulmonary adaptation in disease progression. Biventricular pressure-volume loop data has been recorded previously in rodents^10^, yet this is the first study to use these data for model calibration. Our previous study^4^ showed that model calibration to data from both ventricles reduced parameter and output uncertainty compared to only using RV data. Our study shows that multiorgan models can readily integrate these corresponding data and facilitates more precise model calibration.

Calibrated time course predictions shown in Figure 4(b) show that ventricular and aortic pressures are relatively consistent beat-to-beat. In contrast, ventricular volume measurements vary in magnitude and shape with each cardiac cycle. LV volume measurements recorded by Marquis et al.^18^ were also more variable than the corresponding pressure measurements. Similar variability can be seen in RV volume recordings in the mouse study by Tewari et al.^29,32^. Since there is inherent beat-to-beat variability in these measured quantities, we calibrate our model over multiple heart beats, as shown in Figure 4(a-b).

Uncertainty quantification is a necessary step in the model analysis pipeline. Here we use asymptotic analyses based on frequentist statistical theory^4,18^. Model confidence intervals shown in Figure 4(b) contain most of the LV and RV pressure data. The wider prediction intervals contain most of data, with the largest beat-to-beat variability occurring in the RV volume data. The study by Marquis et al.^18^ also provided output uncertainty in their model predictions using a similar methodology. However, our model confidence intervals are wider and contain a larger proportion of the data.

### 4.3 Simulated ischemia

Myocardial infarction is a precursor to long-term cardiac dysfunction and a risk factor for heart failure. A combined *in-vivo* and *in-silico* analysis provides a potential paradigm for understanding mechanisms of disease progression. Data from all three mice show a reduction in fractional shortening and LV contractile function during ischemia. We simulate impaired LV function by reducing the active force generation by sarcomere shortening. Other authors have considered more sophisticated simulation strategies for ischemia. Witzenburg et al.^34^ separated the LV into infarcted and non-infarcted regions, the latter only contributing to passive LV mechanics. Witzenburg also accounted for compensatory changes in afterload parameters and showed that model predictions matched well with prior experimental (canine) studies. Koopsen et al.^15^ considered a similar, two compartment approach for simulating LV infarction; these authors included biventricular interaction and showed agreement with previously obtained canine data from Lyseggen et al.^17^.

We reduced active force generation to match decreased ejection fraction as measured by echocardiography. LV pressure-volume loops in Figure 5(a) display a rightward shift with ischemia. Shiorua et al.^25^ reported a similar shift in LV pressure-volume loops two weeks and a substantial (nearly 50%) reduction in LV stroke work after mice were subjected to LV ischemia. Simulated RV pressure-volume loops have more subtle changes. Though the RV data during acute LV ischemia do not match our predicted response, both show a leftward shift in the pressure-volume loop, opposite to the LV. Experimentally, Damiano et al.^6^ examined biventricular interaction by excising the sinoatrial node in mongrel dogs and controlling RV pacing. The authors noted that approximately 68% of RV systolic pressure was generated by the LV when ceasing RV pacing in dogs. Our model predicts a small decrease in RV systolic pressure in ischemia but is supported by our experimental measurements of relatively unchanged RV fractional shortening and pressure as measured by catheter. The discrepancy between Damiano’s finding and ours likely results from differences in experimental design (electrical pacing versus ligation), severity of the insult, and species.

Data from both ventricles enhances estimates of ventricular indices, including stroke work (Figure 5(b)). Philip et al.^23^ examined RV pressure-volume loops in mice eight weeks after LV ischemia and saw an increase in RV stroke work relative to the sham animals, which is contrary to our results. Philip et al. attributed this heightened stroke work to the development of pulmonary hypertension after ischemia, the severity of which depends on an increase in pulmonary vascular resistance. Since the increase in resistance is a chronic effect, the RV stroke work likely decreases at the onset of ischemia and then increases as pulmonary pressures rise due to pulmonary vascular remodeling^1^.

Left atrial pressure-volume loops shift upward (Figure 6) with the elevated LV end-diastolic pressure-volume relationship. Bauer et al.^2^ reported an upward shift in bovine atrial pressure-volume loops during acute left anterior coronary artery occlusion. Hanif et al.^13^ reported that mouse models of nonperfused myocardial infarction exhibit left atrial enlargement, atrial cardiomyocyte hypertrophy, and elevated left atrial fibrosis. Elevated left atrial pressure is hypothesized to be a determinant of isolated post-capillary pulmonary hypertension^1^. Philip et al.^23^ showed that LV ischemia in mice increases left atrial wall mass eight weeks after injury. Our model produces elevated pulmonary venous, left atrial, and pericardial pressures during acute LV ischemia, complementing these prior findings.

Our framework includes a model of sarcomere shortening based on a combination of Lumens et al.^16^ and Walmsley et al.^31^. Strain results (Figure 7) confirm equal levels of LV, RV, and S shortening at baseline. In ischemia, there is altered LV shortening and elevated S strain, with no apparent change in RV strain. Dann et al.^7^ compared murine strain magnitudes after LV ischemia and reported significant reductions in LV free-wall shortening within the first seven days post ligation. Clinically, myocardial strain imaging is gaining traction as an indicator of heart function. Hamada-Harimura et al.^12^ reported a strong correlation between RV free-wall longitudinal shortening and adverse cardiac events in acute decompensated heart failure suggesting that incompatible biventricular interactions might be indicative of mortality. The review by Smiseth et al.^26^ identifies several novel uses for strain analysis in LV ischemia, especially during fibrosis and scar development. Further investigations into cardiac wall strain after ischemia are warranted.

Our model framework explicitly models sarcomere length. As LV active force is reduced (i.e., as *γ^*MI*^* approaches zero), end-diastolic volumes and sarcomere lengths reductions in LV active force but change at the prescribed value of *γ^*MI*^* = 0.2. This increase (Figure 8). The shape of the pressure-length curve is maintained for initial “loop” like pattern was observed in Lyseggen et al.^17^, who measured LV long-axis strain in canines during LV ischemia. Using a combination of echocardiography and sonomicrometry, Lyseggen et al. showed that the viable LV myocardial pressure-strain curve switched from counter-clockwise to clockwise after 15 minutes of ischemia. The recent study by Koopsen et al.^15^ reproduced similar plots using a two-compartment model of the ischemic ventricle and simulated the effects of reperfusion that parallel results reported by Lyseggen et al. Our model framework, like Koopsen et al., illustrates the coupled behaviors at both the organ and muscle fiber level during the onset of ischemia, and can be used in future studies related to LV systolic dysfunction.

### 4.4 Limitations

Our study combines a multiscale model of cardiovascular dynamics with pressure-volume loop data from three animals. We plan to use a larger cohort of animals, including both male and female mice, in future studies. We simulate acute LV ischemia but do not account for any acute hemodynamic control mechanisms e.g., the baroreflex. These mechanisms play a role in the long-term homeostasis of the cardiovascular system^34^, but it’s unclear how quickly these response mechanisms act. Future studies across multiple days will require more detailed models of cardiovascular adaptation and remodeling. Lastly, detailed data on the RV response to LV ischemia is necessary. Detailed strain data on biventricular inefficiency and mechanical uncoupling (i.e., a transition from rightward to leftward septal motion) would provide information into the progression of RV dysfunction due to LV dysfunction.

### 4.5 Conclusions

We combine *in-vivo* biventricular pressure-volume loop data with a multiscale computational model of the cardiovascular system. Sensitivity analyses are used to reduce the number of parameters used for inference, and our results show that LV and RV pressure-volume loops can be matched by the model. Our simulations of acute LV ischemia are in line with both recorded RV data and previously published studies documenting the LV’s response. This study displays systems-level hemodynamic changes during the acute stages of myocardial infarction and shows elevated left atrial pressures due to insufficient LV contraction. Our combination of *in-vivo* and *in-silico* techniques provide a framework for understanding the initial effects of LV ischemia and serve as a foundation for an improved understanding of cardiac and vascular remodeling in heart failure.

## COMPETING INTERESTS

The authors declare no conflicts of interest.

## ACKNOWLEDGEMENTS

This work was funded by the National Institutes of Health NIBIB grants R01HL154624 (NCC) and R01 HL147590 (NCC). M.J.C. was supported TL1 TR001415 through the National Center for Research Resources and the National Center for Advancing Translational Sciences, National Institutes of Health. The content is solely the responsibility of the authors and does not necessarily represent the official views of the NIH.

## CITATION DIVERSITY STATEMENT

In agreement with the editorial from the Biomedical Engineering Society (BMES)^24^ on biases in citation practices, we have performed an analysis of the gender and race of our bibliography. This was done manually, though automatic probabilistic tools exist^35^. We recognize existing race and gender biases in citation practices and promote the use of diversity statements like this for encouraging fair gender and racial author inclusion.

Our references contain 21% woman(first)/woman(last), 9% man/woman, 15% woman/man, and 55% man/man. This binary gender categorization is limited in that it cannot account for intersex, non-binary, or transgender people. In addition, our references contain 6% author of color (first)/author of color(last), 3% white author/author of color, 26% author of color/white author, and 65% white author/white author. Our approach to gender and race categorization is limited in that gender and race are assigned by us based on publicly available information and online media. We look forward to future databases that would allow all authors to self-identify race and gender in appropriately anonymized and searchable fashion and new research that enables and supports equitable practices in science.

## Notes

### Competing Interest Statement

The authors have declared no competing interest.

