## Supplementary Material for "Biventricular interaction during acute left ventricular ischemia in mice: a combined in-vivo and in-silico approach"

419 S. Circle View Drive, Suite 6800

Irvine, CA 92697

**Blood volume and pressure estimates**

Nominal parameters are generated using a combination of pressure and volume data or literature. Total blood volume in the mouse is determined by

$$V_{tot}=BW\cdot84.7\frac{\mathrm{kg}}{\mathrm{ml}}$$

(S1)

as postulated by Riches et al.^6^. Bodyweight for mice 1, 2, and 3 were 29, 32.4, and 30.1 (g), respectively, giving $V_{tot}$ values of 2.46, 2.74, and 2.55 ml. Average heart rates for mouse 1, 2, and 3 were 580, 495, and 532 (BPM), respectively. Stroke volume was determined from left (LV) and right ventricle (RV) volume measurements, set to be the max of the two indices. Cardiac output (CO) was calculated as the product of stroke volume and heart rate. The assumed nominal pressures in the other cardiovascular compartment are calculated as a function of the LV and RV pressure data, as detailed in Table 1. The unstressed volumes (i.e., the blood not ejected during a cardiac cycle) and stressed volumes in each compartment are based on previous work^1^ and provided in Table 2. The compartment pressures and stressed volumes are used to construct nominal estimates of vascular resistance and compliance, shown in Table 3.

Table 4 shows the calculated values of the Triseg geometry. Consistent units were necessary for multiscale interactions between model components, hence pressures were converted to KPa using the relation 1 mmHg = 0.133322 KPa, areas were converted from mm^2^ to cm^2^, and volumes were converted from $\mu$l to ml. Outputs from the model are subsequently scaled back for clarity in the results.

**Cardiac Equations**

The sarcomere model is embedded within a cardiac tissue model of atrial dynamics and biventricular interaction (the “TriSeg” model^3^). Changes in blood volume $V\left( t \right)$ ($\mu$l) cause distension in the cardiac chambers, giving rise to the myocardial strain $\varepsilon_{f}$

$$\varepsilon_{f}=\frac{1}{2}\ln\left( \frac{A_{m}}{A_{m,ref}} \right)-\frac{1}{12}z^{2}-0.019z^{4}, z=\frac{3C_{m}V_{wall}}{2A_{m}}.$$

(S2)

Here, $A_{m}$ (mm^2^) is the current mid-wall area of the chamber, $A_{m,ref}$ (mm^2^) is the reference mid-wall area, and $z$ (dimensionless) is a curvature variable related to the ratio of wall volume, $V_{wall}$ (mm^2^), and radius of mid-wall curvature $C_{m}$ (mm^-1^) ^3^. Once $\varepsilon_{f}$ has been calculated and the corresponding $G_{Tot}$ is obtained from the sarcomere model, the mid-wall tension can be calculated as

$$T_{m}=\frac{V_{wall}G_{Tot}}{2A_{m}}\left( 1+\frac{z^{2}}{3}+\frac{z^{4}}{5} \right).$$

(S3)

A balance in axial and radial tensions, $T_{x}$ and $T_{y}$ is enforced across the septal wall

$$\sum_{i=LV,RV,S} T_{x,i}=\sum_{i=LV,RV,S} T_{y,i}=0$$

(S4)

providing two differential algebraic equations ^5^. The cavity tensions are used to calculate the cavity pressures.

The mid-wall volume, $V_{m}$ (mm^3^), mid-wall curvature, $C_{m}$ (mm^-1^), and mid-wall cross-sectional area, $A_{m}$(mm^2^), are the driving variables for cardiac chamber dynamics. In the atria, these are described by

$$V_{m}=V\left( t \right)+\frac{1}{2}V_{wall},$$

(S5)

$$C_{m}=\left( \frac{4\pi}{3V_{m}} \right)^{1/3},$$

(S6)

$$A_{m}=\frac{4\pi}{C_{m}^{2}}.$$

(S7)

Since the LV, RV, and S are mechanically coupled, a separate formulation for $V_{m}$, $C_{m}$, and $A_{m}$ is required (based on the TriSeg model^3^). Utilizing the common radius of mid-wall junction point $y_{m}$ and denoting the maximal axial distance from each chamber wall surface to the origin as $x_{m}$^3^, we get

$$V_{m}=\frac{\pi}{6}x_{m}\left( x_{m}^{2}+3y_{m}^{2} \right)$$

(S8)

$$C_{m}=\frac{2x_{m}}{\left( x_{m}^{2}+y_{m}^{2} \right)},$$

(S9)

$$A_{m}=\pi\left( x_{m}^{2}+y_{m}^{2} \right),$$

(S10)

for the LV, RV, and S. We can also relate $V_{m}$ to the blood volume $V\left( t \right)$ in the chamber

$$V_{m,LV}=-V_{LV}\left( t \right)-\frac{1}{2}V_{wall,LV}-\frac{1}{2}V_{wall,S}+V_{m,S}$$

(S11)

$$V_{m,RV}=-V_{RV}\left( t \right)+\frac{1}{2}V_{wall,LV}+\frac{1}{2}V_{wall,S}+V_{m,S},$$

(S12)

which is updated at each time point.

The atrial transmural pressure is determined from the wall tension and mid-wall curvature

$$p=2T_{m}C_{m}.$$

(S13)

The ventricular mid-wall tension is broken up into the axial ($T_{x}$) and radial ($T_{y}$) tensions based on the geometry of the spherical chambers and the angle of the sphere opening^3^, giving

$$T_{x}=T_{m}\sin\left( \frac{2x_{m}y_{m}}{x_{m}^{2}+y_{m}^{2}} \right), T_{y}=T_{m}\cos\left( \frac{-x_{m}^{2}+y_{m}^{2}}{x_{m}^{2}+y_{m}^{2}} \right).$$

(S14)

These tensions must be balanced across the LV, RV, and septal wall, and serve as the algebraic constraints for the system as described above. The axial tensions in the ventricular cavities are then used to calculate the transmural pressure

$$p=\frac{2T_{x}}{y_{m}}.$$

(S15)

In total, the cardiac chambers and TriSeg model contribute two algebraic constraints ($T_{x}$ and $T_{y}$ being zero), five wall volume parameters ($V_{wall}$), and five reference area parameters ($A_{m,ref}$).

**Table S1.** Formulas for nominal pressure values (presented in mmHg for convenience). Data values are provided in brackets for mouse 1, 2, and 3, respectively.

| **Variable** | **Method/Value** | **Reference** |
| --- | --- | --- |
| $p_{LV,sys}$ | [66.1, 62.3, 55.2] | Data |
| $p_{LV,dias}$ | [5.2, 5.7, 2.1] | Data |
| $p_{RV,sys}$ | [25.0, 19.5, 24.3] | Data |
| $p_{RV,dias}$ | [2.5, 1.5, 1.4] | Data |
| $p_{LA,dias}$ | $0.25\cdot p_{sv,mean}$ | Scaled from normotensive humans ^2^ |
| $p_{RA,dias}$ | $0.25\cdot p_{pv,mean}$ | Scaled from normotensive humans ^2^ |
| $p_{sa,sys}$ | $0.99\cdot p_{LV,sys}$ | Scaled from normotensive humans ^2^ |
| $p_{sa,dias}$ | $0.66\cdot p_{LV,sys}$ | Scaled from normotensive humans ^2^ |
| $p_{sa,mean}$ | $\left( p_{sa,sys}+2p_{sa,dias} \right)/3$ | Clinical definition ^2^ |
| $p_{pa,sys}$ | $0.99\cdot p_{RV,sys}$ | Scaled from normotensive humans ^2^ |
| $p_{pa,dias}$ | $0.32\cdot p_{RV,sys}$ | Scaled from normotensive humans ^2^ |
| $p_{pa,mean}$ | $\left( p_{pa,sys}+2p_{pa,dias} \right)/3$ | Clinical definition ^2^ |
| $p_{sys,cap}$ | $0.26\cdot p_{sa,mean}$ | Scaled from normotensive humans ^2^ |
| $p_{pulm,cap}$ | $0.66\cdot p_{pa,mean}$ | Scaled from normotensive humans ^2^ |
| $p_{sv,mean}$ | $0.26\cdot p_{sa,mean}$ | Scaled from normotensive humans ^2^ |
| $p_{pv,mean}$ | $0.20\cdot p_{pa,mean}$ | Scaled from normotensive humans ^2^ |

**Table S2.** Formulas for nominal unstressed ($V_{i,un}$) and stressed volume ($V_{i}$) values (converted from $\mu l$ to ml$\cdot{10}^{-3}$).

| **Variable** | **Method** | **Reference** |
| --- | --- | --- |
| $V_{sa,un}$ | $0.14\cdot V_{tot}$ | Based on ^2^ |
| $V_{sv,un}$ | $0.70\cdot V_{tot}$ | Based on ^2^ |
| $V_{RA,un}$ | $0.01\cdot V_{tot}$ | Based on ^2^ |
| $V_{RV,un}$ | $0.026\cdot V_{tot}$ | Based on ^2^ |
| $V_{pa,un}$ | $0.026\cdot V_{tot}$ | Based on ^2^ |
| $V_{pv,un}$ | $0.062\cdot V_{tot}$ | Based on ^2^ |
| $V_{LA,un}$ | $0.010\cdot V_{tot}$ | Based on ^2^ |
| $V_{LV,un}$ | $0.026\cdot V_{tot}$ | Based on ^2^ |
| $V_{sa}$ | $0.27\cdot V_{sa,un}$ | Based on ^1^ |
| $V_{sv}$ | $0.075\cdot V_{sv,un}$ | Based on ^1^ |
| $V_{RA}$ | $1.0\cdot V_{RA,un}$ | Based on ^1^ |
| $V_{RV}$ | $1.0\cdot V_{RV,un}$ | Based on ^1^ |
| $V_{pa}$ | $0.58\cdot V_{pa,un}$ | Based on ^1^ |
| $V_{pv}$ | $0.11\cdot V_{pv,un}$ | Based on ^1^ |
| $V_{LA}$ | $1.0\cdot V_{LA,un}$ | Based on ^1^ |
| $V_{LV}$ | $1.0\cdot V_{LV,un}$ | Based on ^1^ |
| $V_{0,peri}$ | $0.9\cdot\left( V_{RA}+V_{RV}+V_{LA}+V_{LV} \right)$ | Hand tuned |
| $V_{m,s}$ | $0.0084\cdot V_{tot}$ | Hand tuned |

**Table S3.** Nominal TriSeg geometry parameter values. Wall volumes ($\mathrm{mm}^{3}$) are calculated as the product of excised chamber weight and myocardial density, assumed to be 1.053 g/ml. Reference areas ($\mathrm{mm}^{2}$) are hand tuned initially to provide reasonable chamber volume predictions. All values are converted to cm in the model.

| **Parameter** | **Method/Value** | **Description** | **Reference** |
| --- | --- | --- | --- |
| $V_{wall,LA}$ | [3.16, 2.42, 4.32] | Wall volume of left atrium | Based on^5^ |
| $V_{wall,LV}$ | [64.4, 85.1, 61.1] | Wall volume of left ventricle | Based on^5^ |
| $V_{wall,RA}$ | [2.21, 4.32, 3.32] | Wall volume of right atrium | Based on^5^ |
| $V_{wall,RV}$ | [22.1, 41.2, 18.5] | Wall volume of right ventricle | Based on^5^ |
| $V_{wall,S}$ | [31.2, 42.5, 30.5] | Wall volume of septum | Based on^5^ |
| $A_{m,ref,LA}$ | [15, 20, 20] | Reference mid-wall area of left atrium | Hand tuned |
| $A_{m,ref,LV}$ | [50,70,50] | Reference mid-wall area of left ventricle | Hand tuned |
| $A_{m,ref,RA}$ | [15, 20, 20] | Reference mid-wall area of right atrium | Hand tuned |
| $A_{m,ref,RV}$ | [55, 80, 50] | Reference mid-wall area of right ventricle | Hand tuned |
| $A_{m,ref,S}$ | [25, 35, 35] | Reference mid-wall area of septum | Hand tuned |

**Table S4.** Formulas for nominal hemodynamic resistances ($R$, KPa$\cdot$s/ml) and compliances ($C$, ml/KPa).

| **Parameter** | **Method/Value** | **Description** | **Reference** |
| --- | --- | --- | --- |
| $R_{m,val}$ | $0.1/CO$ | Mitral valve resistance | Poiseuille/Ohm’s Law |
| $R_{a,val}$ | $0.75/CO$ | Aortic valve resistance | Poiseuille/Ohm’s Law |
| $R_{t,val}$ | $0.1/CO$ | Tricuspid valve resistance | Poiseuille/Ohm’s Law |
| $R_{p,val}$ | $0.75/CO$ | Pulmonic valve resistance | Poiseuille/Ohm’s Law |
| $R_{vc}$ | $\frac{p_{sv,mean} -p_{ra,dias}}{CO}$ | Vena Cava resistance | Poiseuille/Ohm’s Law |
| $R_{pv}$ | $\frac{p_{pv,mean} -p_{la,dias}}{CO}$ | Pulmonary venous resistance | Poiseuille/Ohm’s Law |
| $R_{sys}$ | $\frac{p_{sa,mean} -p_{sys,cap}}{CO}$ | Systemic vascular resistance | Poiseuille/Ohm’s Law |
| $R_{pulm}$ | $\frac{p_{pa,mean} -p_{pulm,cap}}{CO}$ | Pulmonary vascular resistance | Poiseuille/Ohm’s Law |
| $C_{sa}$ | $\frac{V_{sa}}{p_{sa,sys}}$ | Systemic arterial compliance | Based on ^4^ |
| $C_{sv}$ | $\frac{V_{sv}}{p_{sv,mean}}$ | Systemic venous compliance | Based on ^4^ |
| $C_{pa}$ | $\frac{V_{pa}}{p_{pa,sys}}$ | Pulmonary arterial compliance | Based on ^4^ |
| $C_{pv}$ | $\frac{V_{pv}}{p_{pv,mean}}$ | Pulmonary venous compliance | Based on ^4^ |
